# MiR-218: A Molecular Switch and Potential Biomarker of Susceptibility to Stress

**DOI:** 10.1101/589325

**Authors:** Angélica Torres-Berrío, Dominique Nouel, Santiago Cuesta, Eric M. Parise, José María Restrepo-Lozano, Pier Larochelle, Eric J. Nestler, Cecilia Flores

**Author notes:** To whom correspondence may be addressed: Cecilia Flores, Douglas Mental Health University Institute, Department of Psychiatry, McGill University, 6875 Boulevard LaSalle, Montreal, Quebec, H4H 1R3.

## Abstract

Low miR-218 expression in the medial prefrontal cortex (mPFC) is a consistent trait of depression. Here we assessed whether miR-218 in the mPFC confers resilience or susceptibility to depression-like behaviors in adult mice, using the chronic social defeat stress (CSDS) model of depression. We also investigated whether stress-induced variations of miR-218 expression in the mPFC can be detected in blood. We find that downregulation of miR-218 in the mPFC increases susceptibility to a single session of social defeat, whereas overexpression of miR-218 selectively in mPFC pyramidal neurons promotes resilience to CSDS and prevents stress-induced morphological alterations to those neurons. After CSDS, susceptible mice have low levels of miR-218 in the blood as compared to control or resilient groups. We show further that up-and downregulation of miR-218 levels specifically in the mPFC correlates with miR-218 expression in blood. Our results suggest that miR-218 in the adult mPFC might function as a molecular switch that determines susceptibility versus resilience to chronic stress, and that stress-induced variations in mPFC levels of miR-218 could be detected in blood. We propose that blood expression of miR-218 might serve as potential readout of vulnerability to stress and as a proxy of mPFC function.

## INTRODUCTION

Accumulating evidence demonstrates that microRNAs (miRNAs) are important molecular links between environmental risk factors and psychopathology. Altered brain expression of miRNAs has been associated with major depressive disorder (MDD) in humans^1, 2^. These alterations, which localize to brain structures involved in mood regulation, including the prefrontal cortex (PFC), have been observed in humans and rodents exposed to chronic stress^3–7^ and can comprise single miRNAs or clusters of miRNAs^8^. Genome-wide miRNA expression approaches have identified reduced levels of miRNAs involved in neurotransmission, synaptic plasticity and gene regulation in the PFC of antidepressant-free MDD subjects who died by suicide in comparison to non-psychiatric control subjects^4, 6, 7, 9, 10^. In addition, rodents exposed to chronic stress or chronic injection of the stress-related hormone corticosterone display depression-like behaviors and exhibit differential expression of several miRNAs, including miR-214-3p, miR-124-3p and miR-218, in the medial PFC (mPFC)^7, 11–13^.

The diagnosis of MDD relies primarily on the presence of symptoms such as constant feelings of sadness, lack of interest or pleasure, social isolation/avoidance or suicidal ideation. However, given its heterogeneity and the lack of objective biological indicators, the treatment of MDD is still suboptimal for a large percentage of patients^14^. One important feature of miRNAs is that they can be measured in peripheral fluids, including blood, saliva or urine, and their expression could potentially represent a signature of alterations occurring in the central nervous system^15, 16^. Indeed, miRNAs are being recognized as potential diagnostic biomarkers of disease and mediators of antidepressant response to pharmacological and behavioral interventions^2, 15^. Evidence shows that subjects with MDD display reduced levels of miR-1202 and miR-135 in plasma, and those levels can be restored by selective serotonin reuptake inhibitors or cognitive behavioral therapy^4, 5, 17^. Notably, the expression of miR-1202 and miR-135 is altered in postmortem brain tissue of individuals who had been diagnosed with MDD by the time of death. These correlational studies suggest that circulating levels of both miRNAs might reflect alterations occurring in the brain^4, 5^. Other miRNAs have also been reported to be differentially expressed in plasma or peripheral blood mononuclear cells obtained from MDD patients before and after antidepressant treatment, using microarray and small RNA sequencing^17–19^. To date, however, there is still a lack of information about whether and how miRNA alterations in the brain result in measurable changes in their expression in blood and whether those miRNAs are functionally implicated in the vulnerability to MDD.

We previously reported that the miRNA miR-218 is significantly reduced (∼50%) in the PFC of two separate and independent cohorts of non-medicated depressed individuals who died by suicide in comparison to psychiatrically-healthy sudden death controls^7^. We obtained similar findings in mPFC tissue from adult mice exposed to chronic social defeat stress (CSDS), a validated model of depression-like behaviors^20, 21^. Specifically, we showed that mice that developed susceptibility to CSDS exhibit decreased expression (50%) of miR-218 in the prelimbic (PrL) and infralimbic (IL) subregions of the mPFC, in comparison to control and resilient mice^7^. Concordant with the fact that miR-218 is a repressor of the guidance cue receptor gene, *DCC*, we found *DCC* mRNA expression in the PFC to be increased (∼50%) in both depressed suicide individuals and mice susceptible to CSDS^7, 22^. Remarkably, a recent small RNA sequencing study reported miR-218 as one of the miRNAs differentially downregulated in blood obtained from individuals diagnosed with MDD^23^. Together, these findings raise the possibility that variations in miR-218 levels in the mPFC may determine vulnerability to the effects of stress on depression-like behaviors and that these alterations could potentially be observed in blood.

In this study, we used CSDS in rodents, viral infection, and antagomiRs, inhibitors of miRNAs, to investigate this question. We also used mouse blood samples to determine whether stress-induced changes in miR-218 levels in the mPFC can be readily detected peripherally.

## METHODS AND MATERIALS

### Animals

Experimental procedures were performed in accordance with the guidelines of the Canadian Council of Animal Care and approved by the McGill University and Douglas Hospital Animal Care Committee. All mice used in these studies were obtained from Charles River Canada and maintained on a 12h light-dark cycle (light on at 8:00h) with ad libitum access to food and water throughout the experiments. All data derived from animal studies were analyzed by an experimenter blind to experimental conditions.

Male C57BL/6 wild-type mice (PD 75±15) served as experimental subjects in the chronic social defeat stress and the single session of defeat paradigms. Male mice were housed in pairs prior exposure to the stress procedures and single-housed at the completion of the last defeat session and before the social interaction test (SIT). Animals were randomized by cage prior to exposure to stress or before stereotaxic surgeries (i.e. each cage was assigned to each experimental condition or treatment: control or manipulation). The order of the animals was further randomized prior to behavioral tests.

Male CD-1 retired breeder mice (≥3 months old) previously screened for aggressive behavior^7^ were used as social aggressors. CD-1 mice were single-house throughout the study.

### Social Defeat Stress Paradigms

Chronic social defeat stress (CSDS): The CSDS was performed as in^7, 21^. Briefly, each adult male C57BL/6 experimental mouse was exposed to 5 min of physical aggression by a male CD-1 mouse. At the completion of the session, C57BL/6 experimental and CD-1 mice were housed overnight in a 2-compartment rat cage and separated by a transparent divider to provide sensory, but not physical, contact. The procedure was repeated for a total of 10 consecutive days, in which C57BL/6 experimental mice faced a new aggressor every day. Control C57BL/6 mice were housed in similar 2-compartment rat cages with a different littermate every day. The CSDS protocol was conducted during the light cycle, between 11:00 and 14:00.

Single social defeat (SSD): The SSD paradigm consisted of a unique session of social defeat, as described above with minor changes^24^. In brief, adult C57BL/6 experimental mice were exposed to 5 min of physical aggression by a novel CD-1 mouse, and then housed with the same aggressor CD-1 mouse in a 2-compartment rat cage during 15 min to provide psychological stress. After the SSD session, C57BL/6 experimental mice were single-housed 24h prior to the SIT.

Social interaction test (SIT): Twenty-four hr after the last session of CSDS or SSD, C57BL/6 experimental mice were assessed in the SIT as before^7^. This test consisted of 2 sessions in which defeated and control mice explored a squared-arena (42cm × 42cm) in the absence or presence of a novel aggressor CD-1 mouse (social target) for a period of 2.5 min each session. In the first session, an empty wire mesh enclosure (10 cm (w) × 6.5 cm (d) × 42 cm (h)) was located against one of the walls of the arena to assess baseline exploration. In the second session, an unfamiliar CD-1 aggressor was placed inside the wire mesh enclosure. The area that surrounded the enclosure was designated as the social interaction zone (14 cm × 9 cm), whereas the corner area of the wall opposite to the enclosure were designated as corners (9 cm × 9 cm) and represented the farthest point from the social interaction zone. The time (in seconds) of interaction with the social target was estimated during both sessions of the test. The social interaction ratio [time of interaction with social target present/the time in interaction zone with social target absent] was estimated to classify mice as susceptible (ratio<1) and resilient (ratio≥1) as in^7, 21^.

Forced swim test: Immobility was assessed in the forced swim test as described previously^20^. Each mouse was placed in a plexiglass beaker (30 cm high, 15 cm in diameter) for a 5-min session. The beaker was filled with tap water at 23°C and 15 cm in depth. Animal behavior was recorded with video camera for offline manual analysis by an expert observer who was blind to the experimental conditions and registered the duration of immobility during the 5 min of the test. The percentage of immobility time was estimated as the total time of immobility / 5 min*100).

### AntagomiR

A locked Nucleic Acid (LNA) oligonucleotide with a sequence targeting miR-218 (Ant-miR-218) was used to downregulate the expression of miR-218 *in vivo* (Exiqon, Vedbaek, Denmark, Supplementary Table 1). A scrambled LNA oligonucleotide sequence was used as control (Ant-scrambled). Ant-miR-218 or Ant-scrambled were dissolved in sterile PBS (Sigma-Aldrich, Oakville, ON, Canada) at a final concentration of 0.3 mM, as indicated by^25, 26^. We confirmed the efficacy and specificity of Ant-miR-218 (Supplementary Figure 1) by quantitative PCR as in^26^.

### Viral construct

Viral constructs were obtained from Vector BioLabs (Malvern, PA, USA, Supplementary Table 1). Adeno-Associated Virus (AAV8) expressing a pri-mmu-miR-218-GFP fusion protein under the control of the calcium/calmodulin kinase II alpha (CaMKIIα) promoter (miR-218-OE) was used to overexpress miR-218 in pyramidal neurons selectively. A scrambled construct fused to CaMKIIα-GFP was used as control virus (Scrambled).

### Stereotaxic Surgery

All surgeries were performed under aseptic conditions as described in^7^. Adult male C57BL/6 wild-type mice were deeply anesthetized with Isoflurane (5% for induction and 2% for maintenance) and placed in a stereotaxic apparatus. Bilateral microinfusions were made using stainless-steel infusion cannulae (33 gauge) into the mouse mPFC, including PrL and IL subregions, at the following coordinates: +2 mm (A/P), ±0.5 mm (M/L), and −2.7 mm (D/V) relative to Bregma. For miR-218 overexpression experiments, a total volume of 0.75 µl of miR-218-OE (6.9 × 10^13^ genome copies per ml) or Scrambled virus was delivered on each hemisphere over an 8 min period. The infusion cannulae were left inside the brain area during a 6 min pause to prevent virus reuptake. Mice recovered for 21 days before CSDS to maximize peak of expression of the viral constructs. At the completion of the behavioral testing, experimental mice were euthanized for neuroanatomy and gene expression experiments. For miR-218 downregulation experiments, a total volume of 0.5 µl of Ant-miR-218 or Ant-scrambled was infused over a 7 min period. Mice recovered for 7 days prior to exposure to SSD.

### Awake Intranasal Administration of Ant-miR-218 or Ant-Scrambled

Adult male mice were held by the skin of their necks in the palm of the hand to minimize movement as indicated by^27^. A total of 5 µL of Ant-miR-218 or Ant-Scrambled were administered per nostril with a 10XL pipette tip (VWR, Ville Mont-Royal, Quebec, Canada) attached to a P10 micropipette (Corning, Corning, NY, USA). Mice were placed in their home-cage to monitor free movement after the infusion.

### Tissue Dissection and RNA Extraction

Mice that received miR-218-OE virus or antagomiR or that were exposed to CSDS were euthanized by rapid decapitation. Brains were removed, and flash frozen with 2-methylbutane (Sigma-Aldrich, Oakville, ON, Canada) chilled in dry ice. Bilateral punches of the pregenual mPFC, including PrL and IL subregions, were taken from 1 mm coronal sections starting on plate 15 of the Paxinos & Franklin mouse atlas^28^ as previously^7^. Brain punches obtained from the cerebellum correspond to plates 85-88 of the Paxinos & Franklin mouse atlas^28^. Total RNA and miRNA fractions were isolated from the mouse frozen tissue with the miRNeasy Micro Kit protocol (Qiagen, Toronto, ON, Canada). All RNA samples were determined to have 260/280 and 260/230 values ≥1.8, using the Nanodrop 1000 system (Thermo Scientific, Toronto, ON, Canada).

### Blood collection and RNA Extraction

Mice were decapitated, as described above, and blood from trunk was rapidly collected in BD vacutainer tubes containing EDTA as anticoagulant (VWR, Ville Mont-Royal, Quebec, Canada). No cervical dislocation was applied prior to decapitation to maximize the amount of blood collected as approved by our Institutional Animal Care and use Committee. Total RNA from whole blood, including miRNA fraction, was isolated using the Total RNA Purification Kit protocol (Norgen Biotek Corp, Thorold, ON, Canada), according to the manufacturer’s instructions (with minor modifications). All RNA samples were determined to have 260/280 and 260/230 values ≥1.8, using the Nanodrop 1000 system (Thermo Scientific, Toronto, ON, Canada).

### Quantitative Real Time PCR for Mouse Brain Tissue and Whole Blood

Reverse transcription for miR-218 was performed using the TaqMan MicroRNA Reverse Transcription Kit and TaqMan probes (Applied Biosystems, QC, Canada) as in^7^. Real time PCR was run in technical triplicates with an Applied Biosystems QuantStudio RT PCR system (Applied Biosystems, QC, Canada). The small nucleolar RNA (snoRNA) RNU6B and the *C. elegans* specific miRNA, cel-miR-39, were used as endogenous controls of miRNA measures. Expression levels of miR-218 were calculated using the Absolute Quantitation (AQ) standard curve method and normalized by endogenous controls of miRNA as it is recommended for eliminating differences due to sampling as quality of RNA. The expression of miR-218 was relativized to (i.e., normalized to) the control group and presented as fold change.

### Neuroanatomical Experiments with Mouse Brain Tissue

#### In Situ Hybridization

Mice were perfused intracardially with 50 ml of 0.9% saline, followed by 75 ml of ice-cold fixative solution (2% PFA in PBS; pH 7.4). Brains were removed and rapidly frozen with cold 2-methylbutane (Fisher Scientific, Hampton, USA). Coronal sections of the pregenual mPFC were obtained at 12 μm using a cryostat (Leica CM3050 S, Concord, Ontario, Canada) and mounted onto superfrost slides for endogenous peroxidase inactivation with 0.3% H_2_O_2_. Slices were then permeabilized with Proteinase K solution and underwent acetylation with triethanolamine and acetic anhydride. Sense and antisense 5’ digoxigenin-labeled LNA probes against miR-218 (Exiqon, Vedbaek, Denmark; Table S2) were then hybridized to the slices for 14 hr at 60°C. After hybridization, brain sections were stringently washed in saline-sodium citrate (SSC) and 50% formamide for 30 min at 60°C, then treated with RNAse A for 30 hr at 37°C and washed in decreasing concentrations of SSC. mPFC tissue was incubated overnight at 4°C with an anti-DIG antibody coupled to horseradish peroxidase (Roche, Mississauga, ON, Canada) and mouse monoclonal anti-SMI-32 antibody (1:1000 dilution, Previously Covance, Burlington, NC, USA, Cat. #14941802). The anti-SMI-32 antibody was raised against a non-phosphorylated epitope in neurofilament H (NF-H). The non-phosphorylated NF-H is known to be expressed by a subpopulation of pyramidal neurons in layers II, III and V of the rodent and primate cortices^29–32^. To reveal miRNA expression, sections were incubated with tyramide-coupled to Cy3 (Perkin Elmer, Montréal, QC, Canada) and Alexa 488-coupled secondary antibody raised in donkey (Jackson Immunoresearch, Van Allen Way, Carlsbad, CA, USA, Cat. #705-545-003) to reveal SMI-32 immunofluorescence. Sections were rinsed in PBS and coverslipped with DAPI containing media (Vector laboratories, Burlingame, CA, USA).

#### Immunofluorescence

Mice were anesthetized with an overdose of rodent cocktail (a combination of ketamine 50 mg/kg, xylazine 5mg/kg, acepromazine 1mg/kg injected i.p.) and perfused transcardially with 0.9% saline, followed by 4% PFA in PBS (pH 7.4). Brains were immersed in 30% sucrose at 4°C until sunk and flash frozen with cold 2-methylbutane (Fisher Scientific, Hampton, NH, USA). Coronal sections of the pregenual mPFC were obtained at 35 µm using a cryostat (Leica CM3050 S, Concord, ON, Canada). For double-labeled immunofluorescence, endogenous mouse antibodies were blocked using a mouse Ig blocking reagent (Vector Laboratories, Burlingame, USA). mPFC sections were incubated overnight at 4°C with mouse monoclonal anti-SMI-32 antibody (1:1000 dilution, Covance, Burlington, NC, USA, Cat. #14941802) and Goat anti-GFP antibody (1:1000 dilution, Novus Biologicals, Oakville, ON, Canada, Cat. #NB100-1770). Immunostaining was visualized with Alexa 488-conjugated (Jackson Immunoresearch, Van Allen Way, Carlsbad, CA, USA, Cat. #705-545-003), and Alexa Fluor 555-conjugated (Life technologies, Toronto, ON, Canada, Cat. #A21429) secondary antibodies raised in donkey.

#### Stereology

Stereological counts of GFP-positive pyramidal neurons in the pregenual mPFC were performed in a subset of adult wildtype mice infected with miR-218-OE (n=4) or with Scrambled (n=4), as previously^7, 22^. Briefly, the total number of GFP-positive neurons at the site of injection between PrL and IL subregions of the mPFC was evaluated using a stereological fractionator sampling design, with the optical fractionator probe of the StereoInvestigator® software (MicroBrightField, Williston, VT, USA). Regions of interest were delineated according to 5× magnification using a Leica DM4000B microscope (Leica, Concord, ON, Canada). Sections spanning plates 14–18 of the Paxinos & Franklin mouse atlas^28^ were analyzed. Counting was performed at 40× magnification in a 1:4 series. A guard zone of 5μm was used. The coefficient of error (CE) was below 0.1 for all brains. Counts were performed by an expert blinded with respect to experimental conditions.

#### DiL dye staining

Three mice from the four experimental groups were perfused intracardially with saline-heparin solution for 2 min, followed by perfusion with PFA 1.5% in 0.1M phosphate buffer. Whole brains were dissected and post-fixed in PFA 1.5% for 1 hr and then transferred in PBS 0.1M. Coronal sections (200 μm) were collected using a vibratome. Fine powder of DIL crystals (1,1’-Dioctadecyl-3,3,3’,3’-Tetramethylindocarbocyanine Perchlorate, Invitrogen) were suspended in PBS and applied directly to coronal sections for 15 min. Slides were rinsed and kept in PBS at 4°C for 72 hr. Coronal sections were then post-fixed for 30 min in PFA 1.5%. Sections were mounted and cover-slipped with DAPI containing media (Vector laboratories, Burlingame, CA, USA).

#### Analysis of spine density and morphology

Analysis of spine density and morphology was performed on 3-4 coronal sections obtained from control (n=6) or defeated mice (n=6) injected with either miR-218-OE or Scrambled virus as described by^33^. Apical dendrites from PrL and IL subregions of the mPFC were systematically selected for imaging DIL fluorescence, using a confocal microscope (Olympus FV 1200) at 60X immersion objective at 4X zoom and 1024 × 1024 pixels resolution. The Z stack acquisition were performed at 0.3 μm increments. For each group, approximately 8 to 12 of one to four secondary or tertiary dendritic segments from the soma in the apical tree were quantified using NeuronStudio software (http://research.mssm.edu/cnic/tools-ns.html). Dendritic segments from both hemispheres with at least 10 µm of length and that were clearly distinguishable from other segments were included in the analysis. NeuronStudio determines dendrite length semi-automatically and classifies individual spines into thin, mushroom, or stubby according to *(i)* spine aspect ratios, *(ii)* head-to-neck diameter ratios, and *(iii)* head diameters. Thin spines have a neck ratio value (head to neck diameter ratio) less than 1.1 and a length to spine head diameter greater than 2.0. Mushroom spines have a neck ratio value above 1.1 and a spine head diameter equal or greater than 0.3 μm. Stubby spines were discernable by the lack of neck. Each dendritic segment was analyzed separately according to the number of spines and the diameter of the head spines. Spine density was calculated by quantifying the number of spines per 10 μm of dendritic length. The analysis was performed by an experimenter fully blinded across groups.

### Statistical Analysis

Sample sizes (n) are indicated in the figure legends. All values were represented mean ± S.E.M. Statistical analysis was performed using Graphpad Prism 6.0. A significance threshold of α<0.05 was used in all the experiments. Statistical differences between two groups were analyzed with Student’s t-tests with two-tailed analysis. Correlations were calculated using the Pearson correlation coefficient with two-tailed analysis. Otherwise one-way, two-way or three-way ANOVAs were performed, followed by Tukey’s multiple comparison tests. Outliers were screened using the ROUT method. No statistical methods were used to determine the sample sizes, but the number of experimental subjects is similar to sample sizes routinely used in our laboratory and in the field for similar experiments. All data were normally distributed and variance was similar between groups, supporting the use of parametric statistics.

## RESULTS

### Downregulation of miR-218 in the mPFC promotes susceptibility to depression-like behaviors

Reduced expression of miR-218 in the mPFC is associated with depression in humans and susceptibility to CSDS in adult mice^7^. To address whether downregulation of mPFC levels of miR-218 directly causes increased susceptibility to CSDS, we used nucleic acid (LNA)-modified antisense oligonucleotides (antagomiRs) that hybridize miR-218 (Ant-miR-218) with high specificity *in vivo*, leading to its degradation^26^ (Figure 1a; Supplementary Table 1; Supplementary Figure 1). For control micro-infusions, we used LNA-modified antisense oligonucleotides containing a scrambled sequence (Ant-scrambled). One week after infusion, adult mice were exposed to a SSD, which consisted of 5 min of physical aggression by a novel CD-1 mouse, previously screened for aggressive behavior (Figure 1b). Twenty four hr after a SSD, adult mice were tested in the SIT and classified into susceptible and resilient populations according to the social interaction ratio (susceptible <1; resilient >1), as described previously^7^. We chose this paradigm because a single defeat session does not induce social avoidance in adult wild-type mice^24^. As anticipated, a SSD fails to induce social avoidance in mice that received the Ant-scrambled, indicated by the time spent in the interaction zone in the presence of a novel CD1 mouse (social target) (Figure 1c) and the social interaction ratio higher than >1 (Figure 1d). In contrast, a SSD is sufficient to induce high social avoidance in mice injected with Ant-miR-218 in the mPFC (Figure 1c-d), indicating that downregulation of miR-218 in the mPFC induces a pro-susceptible phenotype. Furthermore, miR-218 downregulation in the mPFC increases immobility in the forced swimming test (Figure 1e), another indicator of increased stress susceptibility.

**Figure 1.**
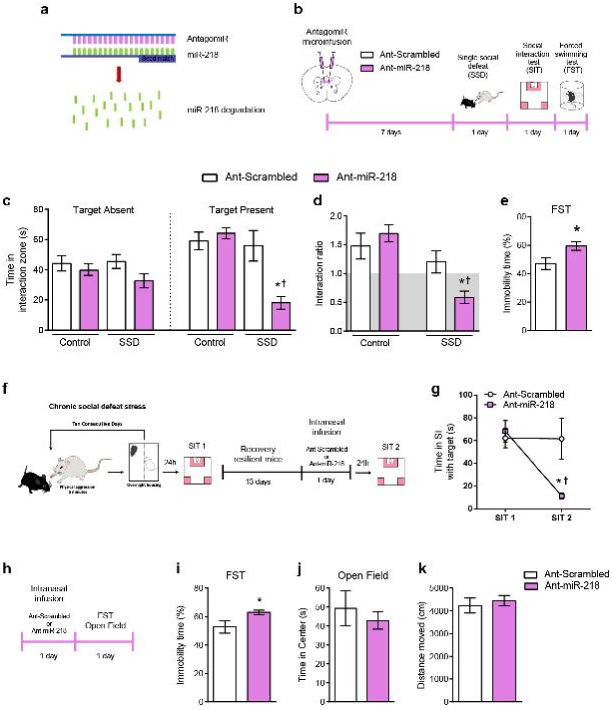
miR-218 downregulation in the mPFC induces depression-like behaviors in mice. **(a)** Schematic illustration of the miR-218 antagomiR (Ant-miR-218). AntagomiRs induce degradation of specific microRNAs *in vivo* upon binding them with high specificity. **(b)** Timeline of experiments. **(c)** Degradation of miR-218 in the mPFC leads to social avoidance following a single session of social defeat (SSD). Time in the interaction zone: Three-way ANOVA: main effect of stress: F(1,58)= 12.40; p=0.001; main effect of target: F(1,58)= 5.022; p=0.029; and main effect of antagomiR: F(1,58)= 10.24; p= 0.002. Significant effect of stress by target by antagomiR interaction: F(1,58)= 4.87; p=0.031. Post hoc Tukey test reveals reduced time in the social interaction zone in SSD-Ant-miR-218 (n=9) during the session with target present as compared to SSD-Ant-scrambled (n=8), and Control-Ant-scrambled (n= 8), *p<0.05; and as compared to Control-Ant-miR-218 (n= 8); ^†^p<0.01. Importantly, Control-Ant-scrambled and Control-Ant-miR-218 groups spend significantly more time exploring the social interaction zone during the session with target present in comparison to target absent; p<0.001. **(d)** Social interaction ratio: Two-way ANOVA: main effect of stress: F_(1,29)_=16.39; p=0.0004; and significant stress by antagomiR interaction: F_(1,29)_=6.03; p=0.02. Post hoc Tukey test shows reduced social interaction ratio in SSD-Ant-miR-218 in comparison to SSD-Ant-scrambled, *p<0.05; and to Control-Ant-scrambled, and Control-Ant-miR-218, ^†^p<0.01. **(e)** Downregulation of miR-218 in the mPFC of adult mice increases percentage of immobility time in the forced swimming test (FST): t_(23)_=2.52; *p= 0.019, different from Ant-scrambled. **(f)** Timeline of CSDS experiment and intranasal infusion of Ant-Scrambled (n=7) or Ant-miR-218 (n=6) in resilient mice. **(g)** Time in the social interaction (SI) zone with Target present before and after intranasal infusion: Two-way ANOVA: main effect of session: F_(1,22)_= 6.098; p= 0.021; main effect of infusion: F_(1,22)_= 3.61; p= 0.07, and significant session by treatment interaction: F_(1,22)_= 5.84; p= 0.024). Post hoc Tukey test shows reduced time in the social interaction zone with the target present in Ant-miR-218 in comparison to Ant-scrambled during SIT 2, *p<0.05; and to Ant-scrambled, and Ant-miR-218 during SIT 1, ^†^p<0.05. **(h)** Timeline of intranasal infusion of Ant-Scrambled or Ant-miR-218 in wild-type mice. **(i)** Intranasal infusion of Ant-miR-218 increases percentage of immobility time in the FST: t_(14)_=2.17; p=0.04. **(j)** Time in center: t_(14)_=0.621; p=0.54. **(k)** Distance moved: t_(14)_=0.55; p=0.59.

### Intranasal delivery of Ant-miR-218 induces susceptibility

Recent evidence demonstrates that intranasal delivery of antagomiRs can successfully alter stress-related behaviors in mice^11^. We therefore explored the possibility that intranasal delivery of Ant-miR-218 could promote depression-like behaviors. First, we exposed adult wild-type mice to CSDS and tested them in the SIT as in^7^. Thirteen days after the first SIT (SIT 1), mice that exhibited a resilient phenotype were infused intra-nasally with either Ant-miR-218 or Ant-Scrambled. Twenty-four h after infusion, mice were tested in a second SIT (SIT 2) to assess for social avoidance (Figure 1f). As shown in Figure 1g, resilient mice infused with Ant-miR-218 spend less time interacting with the social target in the SIT 2 in comparison to SIT 1. This effect was not observed in resilient mice infused with Ant-Scrambled.

To further confirm that reduced miR-218 induces a pro-depressive effect, we infused either Ant-miR-218 or Ant-Scrambled intra-nasally in stress-naïve mice and tested them 24 h later in the forced swimming test (FST) (Figure 1h). We find that intranasal infusion of Ant-miR-218 increases the percentage of immobility time (Figure 1i), again suggesting a pro-susceptible effect. Importantly, two hours after the FST, mice were tested in the open field to assess anxiety-like behaviors and overall locomotion. We did not find significant differences between groups in the time mice spent in the center (Figure 1j) or the total distanced traveled (Figure 1k) in the open field.

### Overexpression of miR-218 in mPFC pyramidal neurons prevents social avoidance induced by CSDS

We processed coronal brain sections derived from stress-naïve adult mice for SMI-32 immunofluorescence and miR-218 in situ hybridization. The SMI-32 antibody was used to label a subset of pyramidal neurons as described previously^7, 22^. We find miR-218 to be expressed in SMI-32-positive pyramidal neurons across all cortical layers of the pregenual mPFC, including cingulate 1, PrL and IL (Figure 2a).

**Figure 2.**
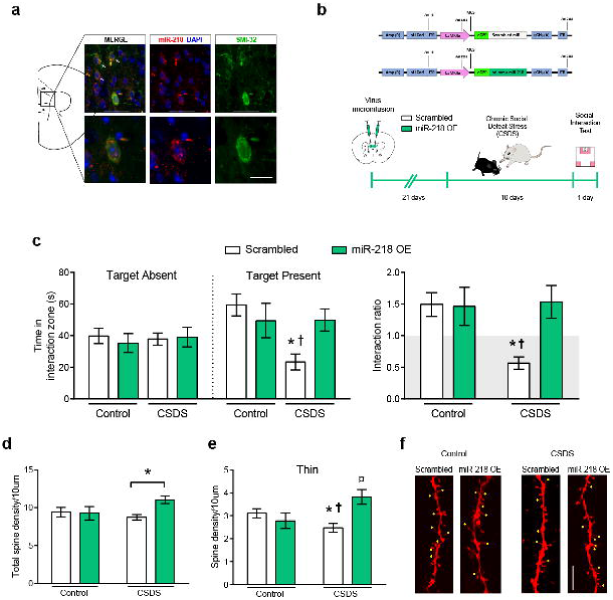
Overexpression of miR-218 in mPFC pyramidal neurons protects against stressinduced CSDS and alterations in spine morphology. **(a)** Representative coronal brain section of a stress-naïve adult mouse showing miR-218 expression in SMI-32-positive mPFC pyramidal neurons (white arrows). Scale bar=10 µm. **(b)** Schematic of the scrambled control (Scrambled) and CaMKIIα-miR-218 overexpression (miR-218 OE) adeno-associated viral (AAV) constructs and timeline of viral infection experiments. **(c)** miR-218 overexpression in mPFC pyramidal neurons prevents social avoidance induced by CSDS and, thereby, promotes resilience. Time in the interaction zone: Three-way ANOVA: No main effect of stress: F_(1,86)_= 2.67; p=0.10; target: F_(1,86)_= 2.05; p=0.15; or virus: F_(1,86)_= 0.80; p=0.37. Significant stress by virus interaction: F_(1,86)_= 4.32; p=0.041. Post hoc Tukey test reveals reduced time in the social interaction zone in CSDS-Scrambled (n=15) during the session with target present in comparison to Control-miR-218 OE (n=11), and CSDS-miR-218 OE (n=12); *p<0.05, and Control-Scrambled (n=9), ^†^p<0.01. Control-Scrambled group spend significantly more time exploring the social interaction zone during the session with target present in comparison to target absent; ^†^p<0.01. Social interaction ratio: Two-way ANOVA: main effect of virus: F_(1,43)_= 4.62; p= 0.03; stress: F_(1,43)_= 3.85; p= 0.056; and significant stress by virus interaction: F_(1,43)_= 5.26; p= 0.026; Tukey test: *p<0.05, different from Control-Scrambled and Control-miR-218 OE; ^†^p<0.01, different from CSDS-miR-218 OE. **(d)** miR-218 overexpression in pyramidal neurons increases dendritic spine density (30%) in the mPFC of socially-defeated mice. Two-way ANOVA: main effect of virus: F_(1,82)_= 3.11; p= 0.08; no significant effect of stress: F_(1,82)_= 0.78; p= 0.37; and significant stress by virus interaction: F_(1,82)_= 3.94; p= 0.05. Post hoc Tukey test: *p<0.05, significantly different from CSDS-Scrambled. **(e)** miR-218 overexpression prevents stress-induced reduction in the density of mPFC thin spines specifically. Two-way ANOVA: main effect of virus: F_(1,82)_= 3.44; p= 0.06; no significant effect of stress: F_(1,82)_= 0.56; p= 0.45; and significant stress by virus interaction: F_(1,82)_= 9.31; p= 0.0031. Post hoc Tukey test: *p<0.05, significantly different from Control-Scrambled, ^†^p<0.01, significantly different from CSDS-miR-218 OE, ¤p<0.01, significantly different from Control-miR-218 OE. **(f)** Representative images of dendritic segments from controls and CSDS-exposed mice injected with Scrambled and miR-218 OE viruses. Yellow dots indicate thin spines. Scale bar, 5μm.

We next assessed whether overexpression of miR-218 in mPFC pyramidal neurons prevents social avoidance after exposure to the CSDS paradigm^7, 21^ (Figure 2b). We microinfused an AAV expressing the primary transcript of miR-218 fused with green fluorescence protein (GFP) under the control of the CaMKIIα promoter (miR-218-OE) bilaterally into the mPFC of adult mice. A scrambled sequence inserted into the AAV-CaMKIIα construct (Scrambled) was used as control virus (Supplementary Table 1; Supplementary Figure 2). Importantly, miR-218-OE and Scrambled constructs are specific to pyramidal neurons and do not compromise neuron integrity or survival (Supplementary Figure 2).

Mice were left in their home cages for 3 weeks following surgery to allow maximal expression of the virus, and then were subjected to a CSDS paradigm, which consisted of 10 consecutive days of physical aggression by a novel CD1 aggressor^7^ (Figure 2b). Twenty-four hr after the last session of defeat, adult mice were tested in the social interaction test^7^. As we anticipated, mice infected with the Scrambled virus and exposed to CSDS exhibit a significant increase in social avoidance: they spend less time interacting with the social target in comparison to non-stressed control mice. By contrast, overexpression of miR-218 selectively in mPFC pyramidal neurons prevents the development of social avoidance in socially-defeated mice, inducing a pro-resilient phenotype (Figure 2c).

We further explored directly the potential role of miR-218 in inducing an antidepressant-like effect. We virally overexpressed miR-218 in the mPFC of wildtype mice that exhibited CSDS-induced susceptibility (Supplementary Figure 2e) and found that miR-218 does not reverse the social avoidance induced by CSDS (Supplementary Figure 2f). This is consistent with the fact that only a small percentage of viral manipulations have successfully reversed CSDS-induced social avoidance^34–37^. Together, our results demonstrate that manipulating miR-218 expression selectively in mPFC pyramidal neurons affect susceptibility to stress-induced depression-like behaviors: while downregulation of miR-218 promotes susceptibility to social stress, miR-218 overexpression prior to stress exposure induces resilience.

### miR-218 increases thin dendritic spines in mice exposed to CSDS

To begin exploring possible mechanisms by which variation of miR-218 in mPFC pyramidal neurons determines vulnerability to depression-like behaviors, we analyzed dendritic segments of mPFC pyramidal neurons in mice subjected to CSDS or control conditions and injected with either miR-218-OE or Scrambled virus. Spine density in dendritic segments of control mice is similar between groups microinjected with miR-218-OE or Scrambled virus (Figure 2d). However, in mice exposed to CSDS, miR-218 overexpression results in a significant increase in spine density (Figure 2d). Because individual spines can be subdivided into mushroom, thin, and stubby, depending on their morphological and functional features^38^, we quantified the density of each type of dendritic spine. Adult mice exposed to CSDS and injected with the Scrambled virus display a significant decrease in the density of thin spines specifically, in comparison to control mice infused with Scrambled virus. This effect is prevented by miR-218 overexpression (Figure 2e). Indeed, socially-defeated mice injected with miR-218-OE have a higher density of thin spines relative to socially-defeated mice injected with Scrambled and control mice infused with miR-218 OE (Figure 2e). It has been suggested that thin spines are highly dynamic and can mature into mushroom spines^39^, and we find a non-significant trend towards an increase in mushroom spine density induced by miR-218 overexpression (Supplementary Figure 2c). There are no significant differences in the density of stubby spines (Supplementary Figure 2c). These results suggest that increased miR-218 in mPFC pyramidal neurons might promote the formation of dendritic spines in animals exposed to CSDS.

### miR-218 can be measured in blood and its levels correlate with social avoidance

A key feature of miRNAs is that they can be measured reliably in several body fluids, including blood, urine, saliva and cerebrospinal fluid^15, 16^. Remarkably, circulating levels of miRNAs can match changes occurring in the brain and can serve as potential biomarkers of psychopathologies^4, 5, 15, 16^. To determine whether stress induces variations of miR-218 in blood, we exposed adult wild-type mice to CSDS. We segregated susceptible and resilient mice and measured the levels of miR-218 in blood 24 hr after the social interaction test (Figure 3a-b). We find that susceptible mice exhibit dramatically (90%) lower levels of miR-218 in blood in comparison to the control and resilient groups (Figure 3c). These results are consistent with our previous report showing reduced mPFC expression of miR-218 of an independent cohort of mice susceptible to CSDS (see Figure 3c - inset)^7^. We next measured the levels of miR-218 in the cerebellum, a brain region highly enriched by miR-218^40^, in the same mice exposed to CSDS and from which we obtained blood samples. As expected, there are not significant differences in miR-218 expression between control, susceptible and resilient mice in the cerebellum (Figure 3d). Together, these findings suggest that stress-induced downregulation of miR-218 in the mPFC might lead to changes of circulating miR-218. Furthermore, there is a significant positive correlation between miR-218 expression in blood and the time spent in the social interaction zone when the social target is present (Figure 3e). Our results indicate a potential role of circulating miR-218 as a readout of stress vulnerability and of mPFC function.

**Figure 3.**
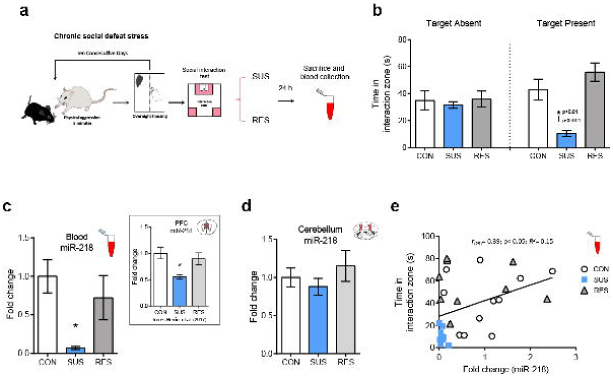
Blood expression of miR-218 correlates with social avoidance. **(a)** Schematic and timeline of CSDS experiment. **(b)** Time in the interaction zone in control (CON; n=11), susceptible (SUS; n=9) and resilient (RES; n=9). Two-way ANOVA: main effect of stress: F_(2,52)_= 8.25; p< 0.001; main effect of session: F_(2,52)_= 0.21; p=0.64; and significant stress by target interaction: F_(2,52)_= 5.58; p<0.01. Post hoc Tukey test: *p<0.01; different from CON; †p<0.001; different from RES, in session with target present. **(c)** Reduced miR-218 levels in blood (∼90%) in SUS: One-way ANOVA: F_(2,26)_= 5.189; p<0.05; Tukey test: *p<0.05; different from CON. Inset represents mean ± SEM of mPFC levels of miR-218 in CON, SUS and RES mice^7^ **(f)** There are no significant differences in the cerebellar levels of miR•218 between CON, SUS and RES mice: One-way ANOVA: F_(2,26)_= 0.84; p=0.44. **(e)** Positive correlation between the levels of miR-218 in blood and the time spent in the interaction zone with the social target present.

### Circulating levels of miR-218 correlate with miR-218 levels in the mPFC

To examine whether circulating levels of miR-218 are a proxy of miR-218 expression in the mPFC, we reduced miR-218 levels in the mPFC and determined whether this manipulation results in downregulation of miR-218 levels in blood. To this end, we micro-infused Ant-miR-218 or Ant-scrambled into the mPFC of adult mice and collected blood samples 10 days after the stereotaxic surgery. We then measured the levels of circulating miR-218 in both groups of mice (Figure 4a). Ant-miR-218 micro-infusion induced a dramatic reduction (80%) in miR-218 expression in the mPFC (Figure 4b) and a strong (∼50%) trend towards a reduction in mR-218 levels in blood (p=0.064). Importantly, there is a significant positive correlation between miR-218 expression in the mPFC and in blood, suggesting that these two variables might be causally related (Figure 4c).

**Figure 4.**
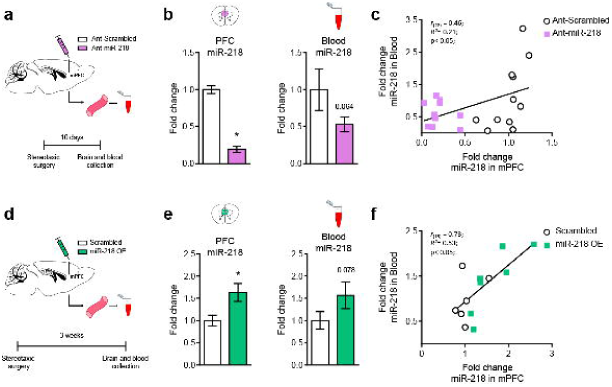
Levels of miR-218 in blood correlate with miR-218 levels in the mPFC. **(a)** Schematic of experimental approach using Ant-miR-218. **(b)** Downregulation of miR-218 in the mPFC tends to decrease miR-218 expression in blood. mPFC: Fold change: t_(20)_= 11.9; p<0.0001. Blood: Fold change: t_(20)_=1.58; p= 0.064. **(c)** Positive correlation between miR-218 levels in mPFC and blood. Ant-scrambled: n= 11; Ant-miR-218: n= 11. **(d)** Schematic of experimental approach using miR-218 OE virus. **(e)** Overexpression of miR-218 in mPFC pyramidal neurons tends to increase miR-218 expression in blood. mPFC: Fold change: t_(11)_= 2.60; p= 0.025. Blood Fold change: t_(11)_= 1.52; p= 0.078. **(f)** Positive correlation between miR-218 levels in mPFC and blood. Scrambled: n= 6; miR-218 OE: n=7.

We next assessed whether overexpressing miR-218 specifically in mPFC pyramidal neurons causes increased circulating levels of miR-218. We microinjected adult mice with either Scrambled or miR-218-OE viruses in the mPFC and then measured blood levels of miR-218 (Figure 4d). Microinjecting the miR-218 OE virus, in comparison to Scrambled infusions, leads to a significant increase (65%) in miR-218 levels in the mPFC and to a similar trend (p=0.078) in blood (Figure 4e), 3 weeks after stereotaxic surgery. Remarkably, there is a high and significant positive correlation between the levels of miR-218 in brain and in blood (Figure 4f). To our knowledge this is the first demonstration that bidirectional manipulations of miR-218 levels in mPFC pyramidal neurons is associated with corresponding changes in miR-218 levels in blood. This finding suggests that the mPFC might be an important source of circulating miR-218^41, 42^.

## DISCUSSION

Depression is the most prevalent psychiatric disorder and represents a high economic burden worldwide^43^. However, the lack of objective biomarkers and the heterogeneity of this condition means that the diagnose relies primarily on self-report of symptoms, increasing the risk of suboptimal treatment^14^. In this context, identifying reliable biomarkers of depression would contribute greatly to the development of novel preventive and treatment strategies. In this study, we provide evidence that miR-218 in the mPFC could be a molecular switch of vulnerability versus resilience to stress-related disorders. Specifically, we report that dampening miR-218 expression in the mPFC promotes stress susceptibility in adult mice and intranasal infusion of a miR-218 antagomiR makes resilient mice susceptible. In contrast, overexpression of miR-218 selectively in mPFC pyramidal neurons protects against the development of depression-like behaviors induced by CSDS, and increases density of thin dendritic spines. In addition, we show that susceptible, but not resilient, mice exposed to CSDS exhibit reduced levels of miR-218 in blood, indicating that circulating miR-218 might serve as a proxy of those changes observed specifically in the mPFC following chronic stress. Indeed, levels of miR-218 in blood correlate positively with social avoidance (Figure 3e), a behavioral trait highly associated with human depression^44^. These findings are strengthened by recent results from a small RNA sequencing study reporting miR-218 as one of the miRNAs differentially downregulated in individuals diagnosed with MDD^23^, further highlighting the promising role of miR-218 as a readout of stress-related disorders.

Determining whether the source of miR-218 in blood is derived at least in part from the mPFC is a provocative and timely question, considering that miRNAs are secreted by cells into the extracellular space^16, 45^. In comparison to other tissues, miR-218 is highly enriched in the brain^46^, mostly in the cortex, and cerebellum, with low but detectable levels in midbrain and striatal regions^26, 40^. Our previous findings demonstrate that miR-218 alterations observed in susceptible mice are mPFC specific: mice susceptible to CSDS exhibit reduced levels of miR-218 in the mPFC, but not in the VTA^7^, a brain region highly involved in susceptibility to CSDS^47^. In this study we further demonstrate that susceptible mice after CSDS do not have changes in miR-218 expression in the cerebellum (Figure 3d), a miR-218-enriched brain region^40^. To our knowledge, this is the first study to report that direct manipulations of the levels of miR-218 in mPFC pyramidal neurons, via viral infection or antagomiR microinfusion, correlate with circulating levels of miR-218, supporting the idea that the mPFC might be an important source of miR-218 in blood.

Circulating miRNAs can be found in either exosomes or in association with proteins or lipoprotein complexes. Exosomes are extracellular microvesicles of 30–120 nm diameter that facilitate direct cell-to-cell interaction by transferring molecular material, including functional proteins, mRNA, and miRNAs, from an active cell to a recipient cell^48, 49^. Exosomes containing miRNAs can be released by neurons and reach different cells, tissues or fluids^48–50^. Future experiments will assess whether mPFC pyramidal neurons release exosomes that contain miR-218, and whether these exosomes reach the blood circulatory system^41, 42^. Indeed, recent evidence demonstrates that motor neurons can release miR-218 to the extracellular space to be taken up by astrocytes^51^. The use of novel technology to trace brain-derived exosomes in combination with bioluminescence imaging *in vivo* or fluorescent microscopy *in vitro* would contribute in this regard^52^. Also, it is important to test the time course of stress-induced changes in circulating levels of miR-218 and whether they correlate with other indicators of stress, including corticosterone. The fact that low levels of circulating miR-218 are observed in mice that become susceptible after CSDS exposure sets the basis for future studies with cohorts of individuals diagnosed with MDD to measure circulating miR-218 before and after antidepressant treatment and determine its association with severity of symptoms, remission or treatment resistance. Overall, our results raise the possibility that circulating miR-218 may be a promising readout of vulnerability to stress and of mPFC function.

MiRNAs can bind to multiple mRNA targets with high spatial and temporal specificity^53^. This regulatory function might be crucial for determining the expression of specific genes in response to environmental challenges. We previously demonstrated that miR-218 represses gene expression of the guidance cue receptor, *DCC*, in human neuroblastoma cell lines^7^. DCC receptors play an important role in the development of the mPFC^54^ and promote synapse formation by allowing synaptic assembly^55^. We also showed that the levels of *DCC* mRNA are significantly increased in the PFC of adult non-medicated MDD subjects who died by suicide, while levels of miR-218 are decreased. This finding suggests that reduced levels of miR-218 in PFC pyramidal neurons renders individuals more susceptible to the deleterious effects of environmental insults in part by increasing the expression of *DCC* mRNA and protein^7^. Indeed, adult mice with selective deletion of *Dcc* in pyramidal neurons of the mPFC exhibit enhanced resilience to CSDS-induced social avoidance and anhedonia^7^. Overexpression of miR-218 in pyramidal neurons similarly protects against the effects of CSDS. The striking similarity in the behavioral effects caused by inducing either miR-218 overexpression or *Dcc* deletion selectively in mPFC pyramidal neurons supports our idea that altered balance of miR-218 and *DCC* is a critical determinant of MDD.

In the brain, miRNAs can control the morphology of pre-and postsynaptic components of synaptic transmission by regulating local translation of dendritic mRNAs^53, 56^. This process depends on neuronal activity and plasticity-related cues that promote miRNA maturation at dendritic sites^53^. Animals susceptible to chronic stress exhibit reduced numbers of dendritic spines and reduced neuronal excitability in mPFC pyramidal neurons^57, 58^. Indeed, rodents exposed to chronic stress display decreased thin, but not mushroom or stubby, spine density in the mPFC^58^. Our neuroanatomy study shows that adult mice exposed to CSDS exhibit a significant decrease in thin spine density in the mPFC as compared to control groups infused with Scrambled virus and that miR-218 overexpression selectively increases the density of thin spines in mice exposed to CSDS. This finding suggests that miR-218 in the mPFC facilitates the formation of thin dendritic spines on pyramidal neurons under stressful conditions, and that these changes in spine plasticity might be protective against the functional and behavioral alterations induced by chronic stress. This idea is supported by recent evidence showing that miR-218 is enriched in forebrain synaptosomes and neurites of primary cultured neurons^59, 60^ and regulates excitatory synaptic transmission at both single neuron and network levels^60^.

## ACKNOWLEDGMENTS AND DISCLOSURES

This work was funded by the National Institute on Drug Abuse (C.F. Grant number: R01DA037911), the Canadian Institute for Health Research (C.F. Grant Number: MOP-74709), the Natural Science and Engineering Research Council of Canada (C.F. Grant Number: 2982226), and the National Institute of Mental Health (E.J.N. Grant numbers: P50MH096890 and R01MH051399). C.F. is a research scholar of the Fonds de Recherche du Québec - Santé. A.T.B. received the Integrated Program in Neuroscience fellowship. We thank Dr. Giovanni Hernandez for his help with the revision of the manuscript, Dr. Elizabeth Ruiz for technical support and Carlos Torres-Berrío for help in the graphic design. The present study used the services of the Molecular and Cellular Microscopy Platform at the Douglas Mental Health University Institute in Montreal, Canada.

The authors declare no potential conflict of interest.

## CONTRIBUTIONS

A.T.B. and C.F. designed the project. A.T.B performed gene expression experiments with mouse brain tissue and blood, stereotaxic surgeries, viral infections, antagomiR infusions and behavioral experiments. D.N and S.C performed all the neuroanatomical experiments. E.M.P. performed behavioral experiments. J.M.R. performed gene expression experiments with mouse brain tissue. P.L. performed stereotaxic surgeries. E.J.N provided reagents and technical training necessary to perform the research. A.T.B and C.F. wrote the manuscript. C.F. supervised the project. All authors discussed the results and commented and edited the manuscript.

## REFERENCES

1. Roy B, Dwivedi Y. miRNAs As Critical Epigenetic Players in Determining Neurobiological Correlates of Major Depressive Disorder. In: Kim Y-K (ed). Understanding Depression: Volume 1. Biomedical and Neurobiological Background. Springer Singapore: Singapore, 2018, pp 51–69.

2. Tavakolizadeh J, Roshanaei K, Salmaninejad A, Yari R, Nahand JS, Sarkarizi HK et al. MicroRNAs and exosomes in depression: Potential diagnostic biomarkers. Journal of Cellular Biochemistry 2018: n/a–n/a.

3. Dias C, Feng J, Sun H, Shao N-y, Mazei-Robison MS, Damez-Werno D et al. β-catenin mediates behavioral resilience through Dicer1/microRNA regulation. Nature 2014; 516(7529): 51–55.

4. Lopez JP, Lim R, Cruceanu C, Crapper L, Fasano C, Labonte B et al. miR-1202 is a primate-specific and brain-enriched microRNA involved in major depression and antidepressant treatment. Nature Medicine 2014; 20(7): 764–768.

5. Issler O, Haramati S, Paul Evan D, Maeno H, Navon I, Zwang R et al. MicroRNA 135 Is Essential for Chronic Stress Resiliency, Antidepressant Efficacy, and Intact Serotonergic Activity. Neuron 2014; 83(2): 344–360.

6. Roy B, Dunbar M, Shelton RC, Dwivedi Y. Identification of MicroRNA-124-3p as a Putative Epigenetic Signature of Major Depressive Disorder. Neuropsychopharmacology 2017; 42(4): 864–875.

7. Torres-Berrío A, Lopez JP, Bagot RC, Nouel D, Dal Bo G, Cuesta S et al. DCC Confers Susceptibility to Depression-like Behaviors in Humans and Mice and Is Regulated by miR-218. Biological Psychiatry 2017; 81(4): 306–315.

8. Geaghan M, Cairns MJ. MicroRNA and Posttranscriptional Dysregulation in Psychiatry. Biological Psychiatry 2015; 78(4): 231–239.

9. Smalheiser NR, Lugli G, Rizavi HS, Torvik VI, Turecki G, Dwivedi Y. MicroRNA Expression Is Down-Regulated and Reorganized in Prefrontal Cortex of Depressed Suicide Subjects. PLOS ONE 2012; 7(3): e33201.

10. Lopez JP, Fiori LM, Gross JA, Labonte B, Yerko V, Mechawar N et al. Regulatory role of miRNAs in polyamine gene expression in the prefrontal cortex of depressed suicide completers. International Journal of Neuropsychopharmacology 2014; 17(1): 23–32.

11. Deng Z-F, Zheng H-L, Chen J-G, Luo Y, Xu J-F, Zhao G et al. miR-214-3p Targets β-Catenin to Regulate Depressive-like Behaviors Induced by Chronic Social Defeat Stress in Mice. Cerebral Cortex 2018.

12. Roy B, Wang Q, Palkovits M, Faludi G, Dwivedi Y. Altered miRNA expression network in locus coeruleus of depressed suicide subjects. Scientific Reports 2017; 7: 4387.

13. Dwivedi Y, Roy B, Lugli G, Rizavi H, Zhang H, Smalheiser NR. Chronic corticosterone-mediated dysregulation of microRNA network in prefrontal cortex of rats: relevance to depression pathophysiology. Transl Psychiatry 2015; 5: e682.

14. Akil H, Gordon J, Hen R, Javitch J, Mayberg H, McEwen B et al. Treatment resistant depression: A multi-scale, systems biology approach. Neuroscience & Biobehavioral Reviews 2018; 84: 272–288.

15. Belzeaux R, Lin R, Turecki G. Potential Use of MicroRNA for Monitoring Therapeutic Response to Antidepressants. CNS Drugs 2017; 31(4): 253–262.

16. Rao P, Benito E, Fischer A. MicroRNAs as biomarkers for CNS disease. Frontiers in Molecular Neuroscience 2013; 6.

17. Lopez JP, Fiori LM, Cruceanu C, Lin R, Labonte B, Cates HM et al. MicroRNAs 146a/b-5 and 425-3p and 24-3p are markers of antidepressant response and regulate MAPK/Wnt-system genes. 2017; 8: 15497.

18. Fan H-m, Sun X-y, Guo W, Zhong A-f, Niu W, Zhao L et al. Differential expression of microRNA in peripheral blood mononuclear cells as specific biomarker for major depressive disorder patients. Journal of Psychiatric Research 2014; 59: 45–52.

19. Belzeaux R, Bergon A, Jeanjean V, Loriod B, Formisano-Tréziny C, Verrier L et al. Responder and nonresponder patients exhibit different peripheral transcriptional signatures during major depressive episode. Transl Psychiatry 2012; 2(11): e185.

20. Krishnan V, Han M-H, Graham DL, Berton O, Renthal W, Russo SJ et al. Molecular Adaptations Underlying Susceptibility and Resistance to Social Defeat in Brain Reward Regions. Cell 2007; 131(2): 391–404.

21. Golden SA, Covington HE, Berton O, Russo SJ. A standardized protocol for repeated social defeat stress in mice. Nature Protocols 2011; 6(8): 1183–1191.

22. Manitt C, Eng C, Pokinko M, Ryan RT, Torres-Berrio A, Lopez JP et al. dcc orchestrates the development of the prefrontal cortex during adolescence and is altered in psychiatric patients. Transl Psychiatry 2013; 3: e338.

23. Mendes-Silva AP, Diniz BS, Tolentino Araújo GT, de Souza Nicolau E, Pereira KS, Silva Ferreira CM et al. MiRNAs and their Role in the Correlation Between Major Depressive Disorder, Mild Cognitive Impairment and Alzheimer’s Disease. Alzheimer’s & Dementia: The Journal of the Alzheimer’s Association 2017; 13(7): P1017–P1018.

24. Razzoli M, Carboni L, Andreoli M, Ballottari A, Arban R. Different susceptibility to social defeat stress of BalbC and C57BL6/J mice. Behavioural Brain Research 2011; 216(1): 100–108.

25. Jimenez-Mateos EM, Engel T, Merino-Serrais P, McKiernan RC, Tanaka K, Mouri G et al. Silencing microRNA-134 produces neuroprotective and prolonged seizure-suppressive effects. Nat Med 2012; 18(7): 1087–1094.

26. Cuesta S, Restrepo-Lozano JM, Silvestrin S, Nouel D, Torres-Berrío A, Reynolds LM et al. Non-Contingent Exposure to Amphetamine in Adolescence Recruits miR-218 to Regulate Dcc Expression in the VTA. Neuropsychopharmacology 2017.

27. Hanson LR, Fine JM, Svitak AL, Faltesek KA. Intranasal Administration of CNS Therapeutics to Awake Mice. JoVE 2013; (74): e4440.

28. Paxinos G, Franklin K. Paxinos and Franklin’s the Mouse Brain in Stereotaxic Coordinates. Elsevier/Academic Press: Boston, Amsterdam, 2013.

29. Campbell MJ, Morrison JH. Monoclonal antibody to neurofilament protein (SMI-32) labels a subpopulation of pyramidal neurons in the human and monkey neocortex. The Journal of Comparative Neurology 1989; 282(2): 191–205.

30. Voelker CCJ, Garin N, Taylor JSH, Gähwiler BH, Hornung J-P, Molnár Z. Selective Neurofilament (SMI-32, FNP-7 and N200) Expression in Subpopulations of Layer V Pyramidal Neurons In Vivo and In Vitro. Cerebral Cortex 2004; 14(11): 1276–1286.

31. Sternberger LA, Sternberger NH. Monoclonal antibodies distinguish phosphorylated and nonphosphorylated forms of neurofilaments in situ. Proceedings of the National Academy of Sciences of the United States of America 1983; 80(19): 6126–6130.

32. Hof PR, Morrison JH. Neurofilament protein defines regional patterns of cortical organization in the macaque monkey visual system: A quantitative immunohistochemical analysis. The Journal of Comparative Neurology 1995; 352(2): 161–186.

33. Mahmmoud RR, Sase S, Aher YD, Sase A, Gröger M, Mokhtar M et al. Spatial and Working Memory Is Linked to Spine Density and Mushroom Spines. PLOS ONE 2015; 10(10): e0139739.

34. Vialou V, Robison AJ, Laplant QC, Covington HE, 3rd, Dietz DM, Ohnishi YN et al. DeltaFosB in brain reward circuits mediates resilience to stress and antidepressant responses. Nature Neuroscience 2010; 13(6): 745–752.

35. Golden SA, Christoffel DJ, Heshmati M, Hodes GE, Magida J, Davis K et al. Epigenetic regulation of RAC1 induces synaptic remodeling in stress disorders and depression. Nature Medicine 2013; 19: 337.

36. Sun H, Damez-Werno DM, Scobie KN, Shao N-Y, Dias C, Rabkin J et al. ACF chromatin-remodeling complex mediates stress-induced depressive-like behavior. Nat Med 2015; 21(10): 1146–1153.

37. Jiang C, Lin WJ, Sadahiro M, Labonté B, Menard C, Pfau ML et al. VGF function in depression and antidepressant efficacy. Mol Psychiatry 2018; 23(7): 1632–1642.

38. Hering H, Sheng M. Dentritic spines: structure, dynamics and regulation. Nat Rev Neurosci 2001; 2(12): 880–888.

39. Spruston N. Pyramidal neurons: dendritic structure and synaptic integration. Nature Reviews Neuroscience 2008; 9: 206.

40. Bak M, Silahtaroglu A, Møller M, Christensen M, Rath MF, Skryabin B et al. MicroRNA expression in the adult mouse central nervous system. RNA 2008; 14(3): 432–444.

41. Bala S, Csak T, Momen-Heravi F, Lippai D, Kodys K, Catalano D et al. Biodistribution and function of extracellular miRNA-155 in mice. 2015; 5: 10721.

42. Laulagnier K, Javalet C, Hemming FJ, Sadoul R. Purification and Analysis of Exosomes Released by Mature Cortical Neurons Following Synaptic Activation. In: Hill AF (ed). Exosomes and Microvesicles: Methods and Protocols. Springer New York: New York, NY, 2017, pp 129–138.

43. Chisholm D, Sweeny K, Sheehan P, Rasmussen B, Smit F, Cuijpers P et al. Scaling-up treatment of depression and anxiety: a global return on investment analysis. The Lancet Psychiatry 2016; 3(5): 415–424.

44. Krishnan V, Nestler EJ. The molecular neurobiology of depression. Nature 2008; 455(7215): 894–902.

45. Chen RJ, Kelly G, Sengupta A, Heydendael W, Nicholas B, Beltrami S et al. MicroRNAs as biomarkers of resilience or vulnerability to stress. Neuroscience 2015; 305: 36–48.

46. Small EM, Sutherland LB, Rajagopalan KN, Wang S, Olson EN. MicroRNA-218 Regulates Vascular Patterning by Modulation of Slit-Robo Signaling. Circulation Research 2010; 107(11): 1336–1344.

47. Chaudhury D, Walsh JJ, Friedman AK, Juarez B, Ku SM, Koo JW et al. Rapid regulation of depression-related behaviours by control of midbrain dopamine neurons. Nature 2013; 493(7433): 532–536.

48. Chivet M, Hemming F, Pernet-Gallay K, Fraboulet S, Sadoul R. Emerging Role of Neuronal Exosomes in the Central Nervous System. Frontiers in Physiology 2012; 3(145).

49. Valadi H, Ekstrom K, Bossios A, Sjostrand M, Lee JJ, Lotvall JO. Exosome-mediated transfer of mRNAs and microRNAs is a novel mechanism of genetic exchange between cells. Nat Cell Biol 2007; 9(6): 654–659.

50. Korkut C, Li Y, Koles K, Brewer C, Ashley J, Yoshihara M et al. Regulation of Postsynaptic Retrograde Signaling by Presynaptic Exosome Release. Neuron 2013; 77(6): 1039–1046.

51. Jensen LA, Reddy LV, Hoye ML, Miller TM, Richard J-P, Rothstein JD et al. Motor neuron-derived microRNAs cause astrocyte dysfunction in amyotrophic lateral sclerosis. Brain 2018; 141(9): 2561–2575.

52. Arenaccio C, Chiozzini C, Ferrantelli F, Leone P, Olivetta E, Federico M. Exosomes in Therapy: Engineering, Pharmacokinetics and Future Applications. Current Drug Targets 2019; 20(1): 87–95.

53. Sambandan S, Akbalik G, Kochen L, Rinne J, Kahlstatt J, Glock C et al. Activity-dependent spatially localized miRNA maturation in neuronal dendrites. Science 2017; 355(6325): 634–637.

54. Hoops D, Flores C. Making Dopamine Connections in Adolescence. Trends in Neurosciences 2017; 40(12): 709–719.

55. Goldman JS, Ashour MA, Magdesian MH, Tritsch NX, Harris SN, Christofi N. Netrin-1 promotes excitatory synaptogenesis between cortical neurons by initiating synapse assembly. J Neurosci 2013; 33.

56. Schratt GM, Tuebing F, Nigh EA, Kane CG, Sabatini ME, Kiebler M et al. A brain-specific microRNA regulates dendritic spine development. Nature 2006; 439(7074): 283–289.

57. Radley JJ, Rocher AB, Miller M, Janssen WGM, Liston C, Hof PR et al. Repeated Stress Induces Dendritic Spine Loss in the Rat Medial Prefrontal Cortex. Cerebral Cortex 2006; 16(3): 313–320.

58. Bloss EB, Janssen WG, Ohm DT, Yuk FJ, Wadsworth S, Saardi KM et al. Evidence for Reduced Experience-Dependent Dendritic Spine Plasticity in the Aging Prefrontal Cortex. The Journal of Neuroscience 2011; 31(21): 7831–7839.

59. Siegel G, Obernosterer G, Fiore R, Oehmen M, Bicker S, Christensen M et al. A functional screen implicates microRNA-138-dependent regulation of the depalmitoylation enzyme APT1 in dendritic spine morphogenesis. Nat Cell Biol 2009; 11(6): 705–716.

60. Rocchi A, Moretti D, Lignani G, Colombo E, Scholz-Starke J, Baldelli P et al. Neurite-Enriched MicroRNA-218 Stimulates Translation of the GluA2 Subunit and Increases Excitatory Synaptic Strength. Molecular Neurobiology 2019.

