## Supplementary Information for "MiR-218: A Molecular Switch and Potential Biomarker of Susceptibility to Stress"

### Supplementary Figure 1

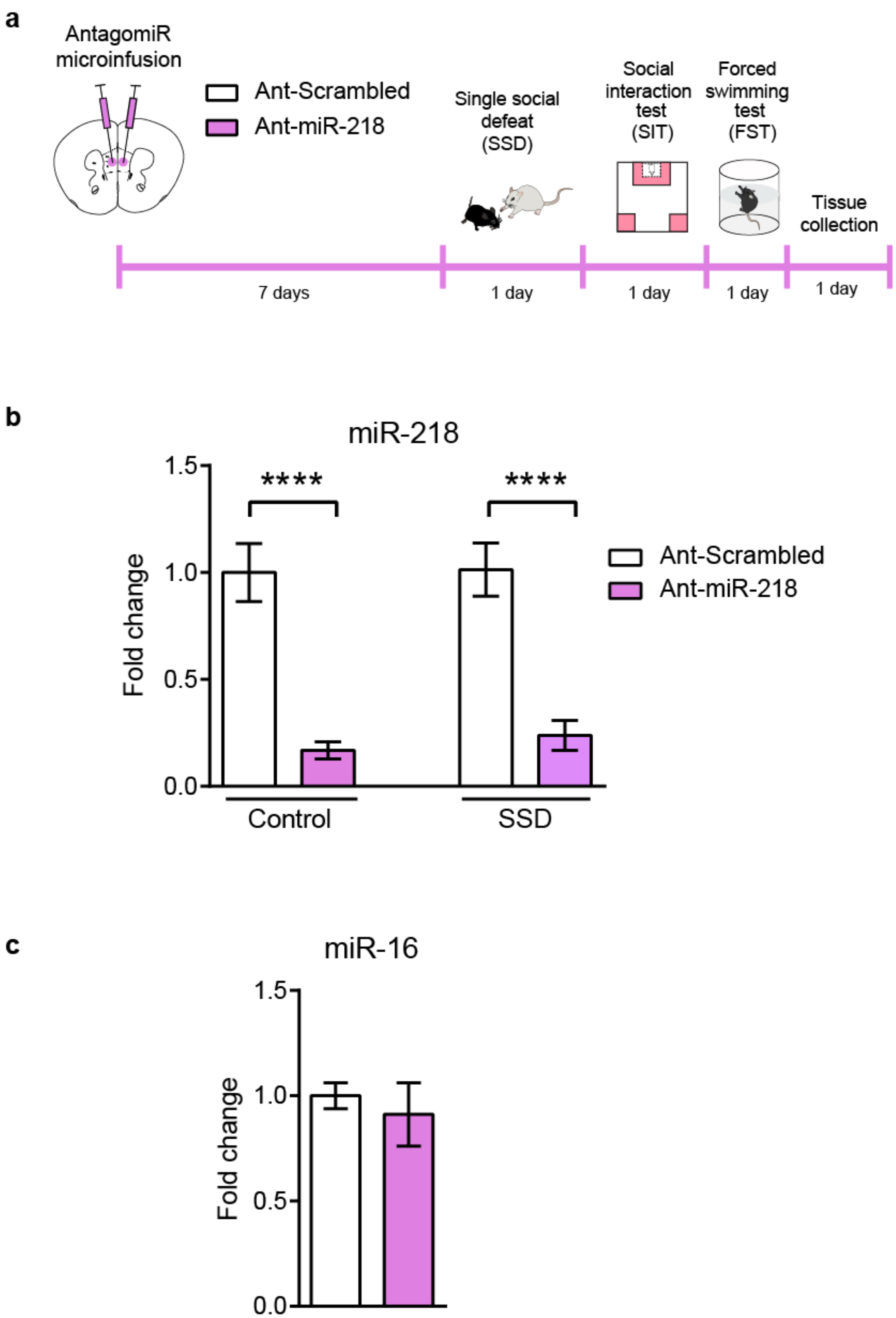

**Supplementary Figure 1. Validation of the antagomiR for miR-218. (a)** Timeline of antagomiR infusion and behavioral experiments included in Figure 1. **(b)** Ant-miR-218 induced a significant reduction of miR-218 in the mPFC of mice exposed to SSD (SSD-Ant-miR-218; n=9) or control (Control-Ant-miR-218; n= 8) in comparison to SSD-Ant-scrambled (n=8), and Control-Ant-scrambled (n= 8), indicating that inhibition of miR-218 by Ant-miR-218 was consistent across animals: Two-way ANOVA: Main effect of antagomiR: $F(_{1,29})= 59.94$; $p<0.0001$; Main effect of group: $F_{(1,29)}= 0.165$; $p=0.69$; antagomiR by group interaction: $F_{(1,29)}= 0.073$; $p=0.78$. Post hoc Tukey test shows reduced levels of miR-218 in mPFC of mice infused with Ant-miR-218 in comparison to Ant-Scrambled, ****$p<0.0001$. **(c)** Adult stress-naïve mice infused with Ant-miR-218 (n=6) and Ant-Scrambled (n=6) exhibited similar levels of miR-16, a miRNA that is not predicted to be affected by Ant-miR-218: $t_{(10)}=0.54$; $p=0.6$.

### Supplementary Figure 2

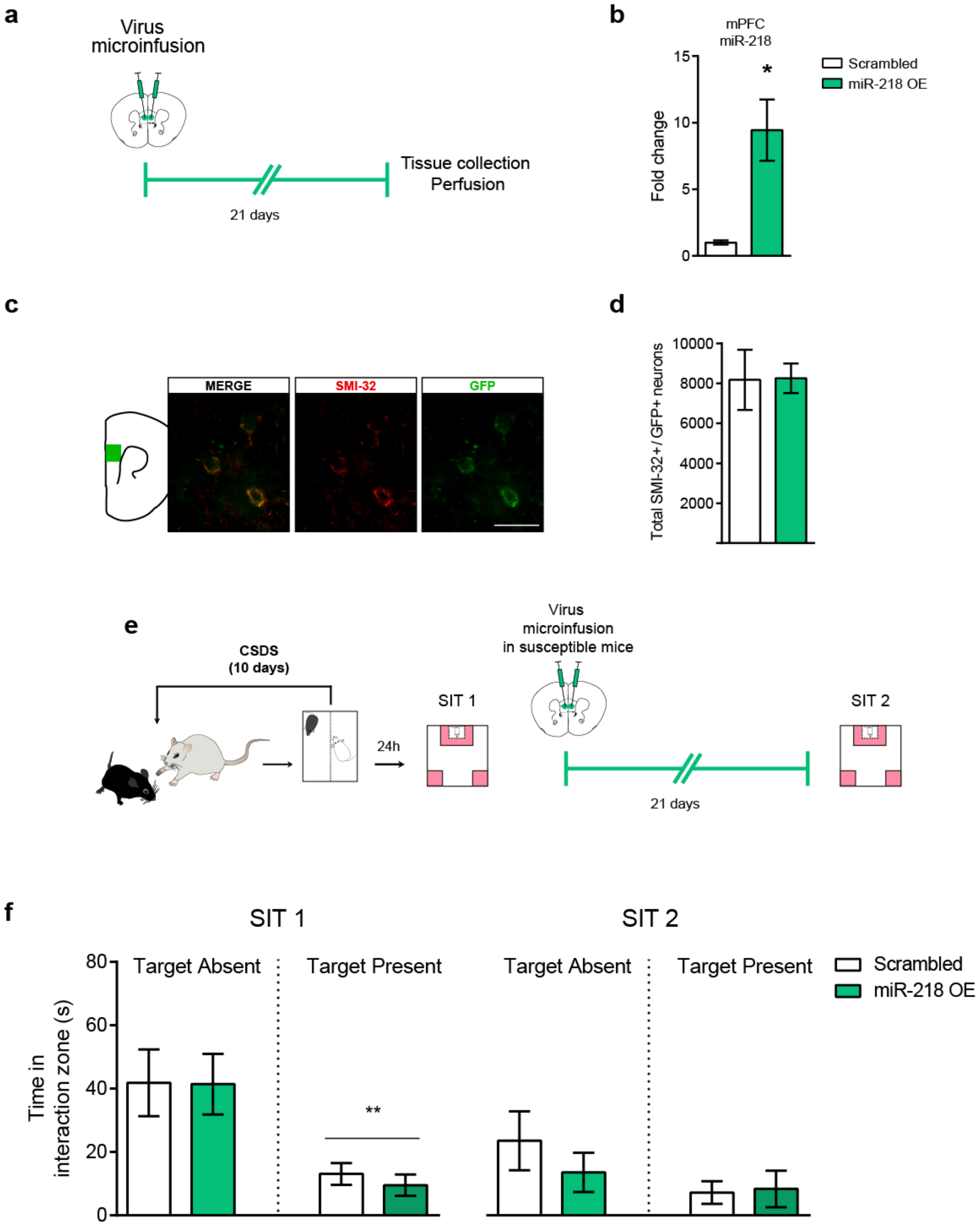

**Supplementary Figure 2. Validation of the miR-218 overexpression (miR-218-OE) virus. (a)** Timeline of miR-218-OE or (Scrambled) AAVs microinfusion in the mPFC. **(b)** miR-218 levels in the mPFC (n=5) of mice injected with miR-218-OE or Scrambled viruses under basal conditions: $t_{(8)}$=3.669; p= 0.0063. **(c)** GPF expression in SMI-32-positive neurons in the pregenual mPFC of adult wild-type mice injected miR-218-OE, confirming specificity pyramidal neurons of our viral construct. Scale bar= 50µm. **(d)** No significant differences in the total number of SMI-32+/GFP infected neurons between groups infected with miR-218-OE or Scrambled viruses: ($t_{(6)}$=0.14; p= 0.89). **(e)** miR-218-OE in the mPFC does not reverse susceptibility to CSDS. Adult male wildtype mice exposed to CSDS were tested in the social interaction test (SIT 1). Mice that exhibited a susceptible phenotype were microinfused with either miR-218-OE (n=4) or Scrambled (n=4) viruses in the mPFC, and were tested for a second social interaction test (SIT 2) twenty-one days after viral infection. **(f)** There are no significant differences between groups in the time spent in the social interaction zone in the absence or presence of a social target in the SIT 1 and SIT 2: Three-way ANOVA: Main effect of target: $F_{(1,24)}$=18.64; p<0.0001; Main effect of SIT: $F_{(1,24)}$= 5.18; p=0.032; No significant effect of Virus: $F_{(1,24)}$=1.04; p=0.316; SIT by target interaction: $F_{(1,24)}$=3.057; P=0.093; SIT by Virus interaction: $F_{(1,24)}$=0.39; p=0.53; Virus by Target interaction: $F_{(1,24)}$=0.36; p=0.55; SIT by Virus by Target: $F_{(1,24)}$= 0.84; p=0.36.

### Supplementary Figure 3

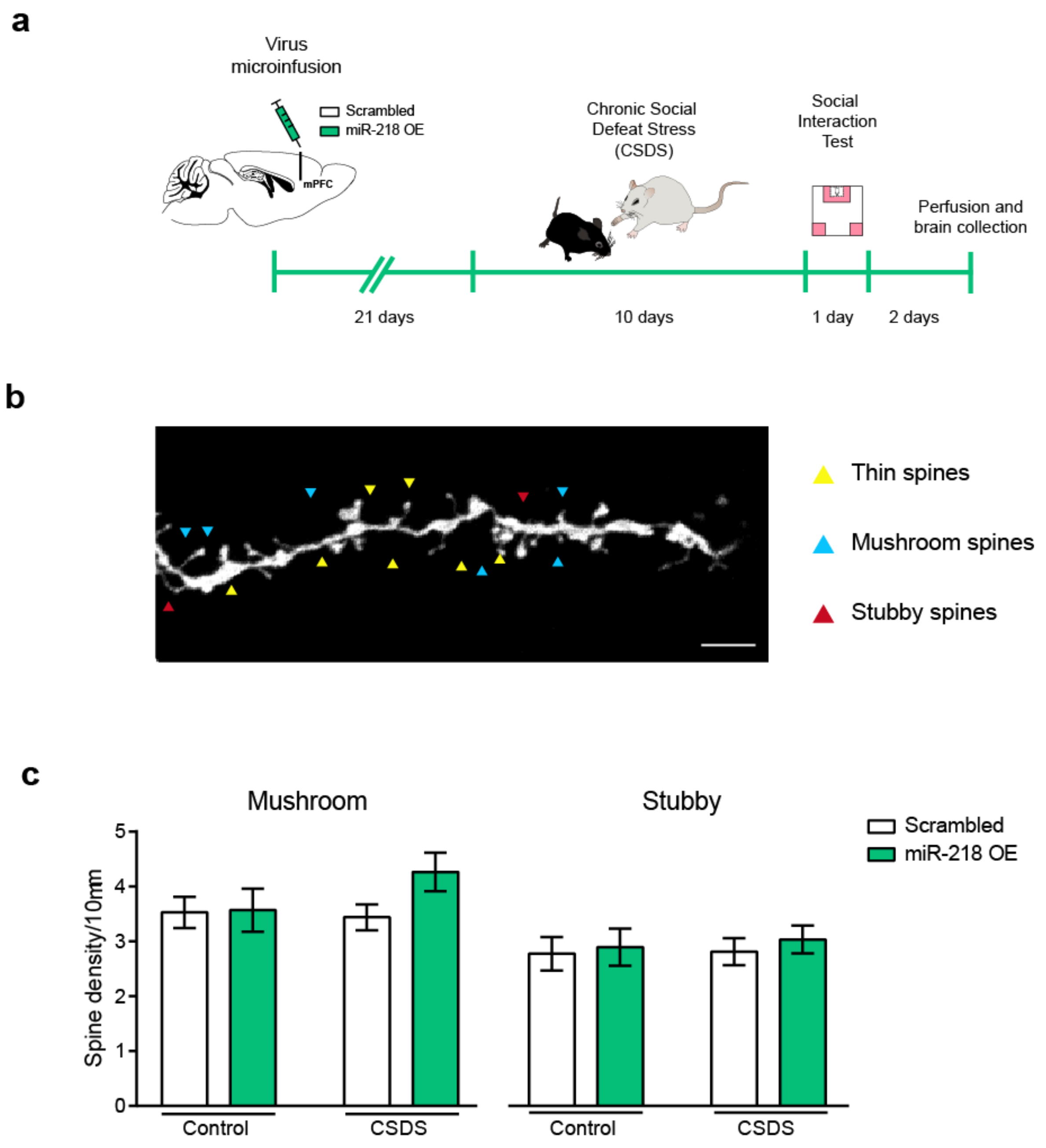

**Supplementary Figure 3. miR-218 overexpression in the mPFC does not change mushroom or stubby spine density. (a)** Timeline of viral infection, behavioral and neuroanatomy experiments. **(b)** Representative high-magnification image of a dendritic segment. Thin, Mushroom and Stubby spine subtypes are indicated by pointing arrows. Scale bar, 5μm. **(c)** Mushroom spines: Two-way ANOVA: virus: $F_{(1,82)}= 1.96$; $p= 0.16$; stress: $F_{(1,82)}= 0.78$; $p= 0.37$; and virus by stress interaction: $F_{(1,82)}= 1.39$; $p= 0.24$. Stubby spines: Two-way ANOVA: virus: $F_{(1,82)}= 0.35$; $p= 0.55$; stress: $F_{(1,82)}= 0.09$; $p= 0.75$; and virus by stress interaction: $F_{(1,82)}= 0.031$; $p= 0.85$.

**Supplementary Table 1.**

Sequence of Adeno-associated viruses and AntagomiRs.

| Gene Target | Accession Number | Sequence | Application |
|---|---|---|---|
| mmu-miR-218-5p | | GUGAUAAUGGAGCGAGAUUUUCUGUU GUGCUUGAUCUAACCAUGUGCUUGCG AGGUAUGAGAAAAACAUGGUUCCGUC AAGCACCAUGGAACGUCACGCAGCUU UCUACA | Virus |
| Scrambled | | AGTCTCCACGCGCAGTACATTTTAGTG AAGCCACAGATGTAAAATGTACTGCGC GTGGAGACC | Virus |
| mmu-miR-218-5p | MIMAT0000663 | UUGUGCUUGAUCUAACCAUGU | AntagomiR |
| Scrambled | | GGTTAGATCAAGCACA | AntagomiR |

**Supplementary Table 2**.

List of antibodies used for immunofluorescence, western blot and in situ hybridization experiments.

| Antigen | Immunogen | Manufacturer | Application | Dilution |
|---|---|---|---|---|
| SMI-32 | Raised against a nonphosphorylated epitope in neurofilament H. Recognizes a subpopulation of pyramidal neurons. | Covance, (Cat. #14941802), monoclonal mouse | ICC | 1:1000 |
| α-DIG-POD | Polyclonal antibody specific to digoxigenin and digoxin. | Roche, (Cat. #11207733910), sheep polyclonal | ISH | 1:2500 |
| α-DIG AP | Polyclonal antibody specific to digoxigenin and digoxin. | Roche, (Cat. #11093274910), sheep polyclonal | ISH | 1:1000 |
